# Host’s Specific SARS-CoV-2 Mutations: Insertion of the Phenylalanine in the NSP6 Linked to the United Kingdom and Premature Termination of the ORF-8 Associated with the European and the United States of America Derived Samples

**DOI:** 10.1101/2020.12.29.424530

**Authors:** Mohammad Khalid, Yousef Al-ebini, David Murphy, Maryam Shoai

**Author notes:** **Corresponding author:** Mohammad Khalid, Department of Pharmaceutics, College of Pharmacy, King Khalid University, Abha-61421, Asir, Saudi Arabia.

## Abstract

The coronavirus belongs to the order Nidovirales, which is known for the longest RNA genome virus. The polymerase enzyme of SARS-CoV-2 has proofreading functions, but still, the RNA viruses have a higher mutation rate than DNA viruses. The mutations in the viral genome provide a replication advantage in any population/geographical location and that may have profound consequences in the outcome and pathogenesis, diagnosis and patient management of the viral infection. In the present study, we have analysed full-length SARS-CoV-2 genome sequences, derived from symptomatic/asymptomatic COVID-19 patients from all six continents to investigate the common mutations globally. Our results revealed that SARS-CoV-2 is mutating independently, we identified total 313 mutations and some (21 mutations) of them are prevailing over time irrespective of geographical location. Another important finding, we are reporting here is, the mutation rate of the virus varies in different geographical locations suggesting the virus is adapting different strategies in the infected populations, having different genetic backgrounds across the globe. We have identified 11085TTT insertion (insertion of the Phenylalanine in NSP6 at position 38) mutation, which is mainly linked to the UK derived SARS-CoV-2 samples, we have also discovered non-sense mutation in ORF-8 after 17 amino acid is linked to the European and the USA derived SARS-CoV-2 samples.

## Introduction

The coronaviruses belong to the order Nidovirales, family Coronavirdiae, and subfamily Coronavirinae (1). The COVID-19 pandemic is caused by the novel coronavirus, named as 2019-nCoV or SARS-CoV-2, belonging to the Betacoronavirus (βCoV) genus of the subfamily Coronavirinae. The SARS-CoV-2 virus contains a positive-sense single-stranded RNA (+ssRNA) genome with a size of around 29.9 kilobases (2). The genomes of coronaviruses are among the longest RNA viruses (3). The reverse transcription of the genome is needed after the viral entry into the host cell for further replication, which is performed by the RNA-dependent RNA polymerase (RdRp) enzyme of the viruses. Although coronavirus’ RdRp have a proofreading function, single-stranded RNA genomes have a higher mutation rate than double-stranded DNA viruses (4). Another cause of mutation of the viral genome is the driving force to evade the defences of the immune system of the host. These factors make mutant variants of various viral proteins, which help the virus in evading the defences. Consequently, the fitting variants prevail in any geographical region/population over time.

The change in nucleotide/amino acid sequences may have the consequences of changes in the viral infectivity, poor efficiency of molecular diagnosis and escape from antibody recognition. Consequently, the failure of the vaccine and the development of drug-resistant viral strains could arise. Furthermore, the mutational changes in viral receptor binding proteins would result in change in tropism that might bring expanded host range or zoonotic transfer.

The mutational changes in the viral genome may lead to sequence changes in untranslated regions (UTRs) as well as changes in amino acid sequences of structural and functional proteins of the viruses. Being the RNA virus, the SARS-CoV-2 is mutating; several rare and common mutations in UTR/ORFs in the genome have been already reported (5-10). The D614G mutation in spike protein has got special attention among the SARS-CoV-2 virologist for becoming dominant around the world (11) and the mutation also correlated with a higher mortality rate in COVID-19 patients (12). It is highly likely that the SARS-CoV-2 may have more mutations like D614G, which may affect the viral infectivity and clinical outcome of the infection drastically that needs to be explored more thoroughly.

Our aim was to analyse the SARS-CoV-2 genome sequences derived from patients of all continents and to investigate whether the virus mutate in similar pattern globally or has specific pattern in any given populations. In the present, study we analysed 15,120 full length SARS-CoV-2 genomes, derived from symptomatic/asymptomatic COVID-19 patients from all six continents, submitted to NCBI Database (13). We further investigated an additional 1,000 SARS-CoV-2 genome sequences, derived from Australia, China, India, KSA, Egypt, Italy, Spain, USA, Mexico and Brazil and submitted to GISAID (14).

## Methods

### SARS-CoV-2 Genome Sequences

In the present study, we analysed 15,120 full-length SARS-CoV-2 genomes (global set), derived from symptomatic/asymptomatic COVID-19 patients. These sequences were submitted to NCBI database from different countries covering all six continents. We downloaded all the SARS-CoV-2 genome sequences available from the NCBI database as on June 25, 2020.

We also investigated the 1,000 more SARS-CoV-2 genome sequences (country set) from symptomatic/asymptomatic COVID-19 patients from 10 countries (Australia, China, India, KSA, Egypt, Italy, Spain, USA, Mexico and Brazil, covering all six continents). Large number of SARS-CoV-2 infection cases have been reported from these countries. The sequences were downloaded from GISAID and 100 full-length SARS-CoV-2 genome sequences were randomly selected from each country.

### Mutational Analysis of the SARS-CoV-2 Genome Sequences

The 15,120 full-length SARS-CoV-2 genome sequences were processed using FASTA_Unique_Sequences_1.0 tool (15), which removed the base-for-base duplicates and that brought us down to the 13,136 full length genome sequences. Next, we used a sequencing tool, Minimap2 (16), which is designed for long-read sequences with a larger number of samples to be aligned. Using Minimap2, once aligned with a reference sequence (accession number NC_045512) (17), the duplicate sequences were removed more aggressively and that gave us 400 genome sequences with mutations (Workflow: Step A). Although the MiniMap2 is suited for longer sequences and large number of samples, it has the disadvantage of showing the most common mutations only and hence we had to utilize another approach to find less common mutations.

To analyse the 1000 full-length SARS-CoV-2 genome sequences, derived from every above-mentioned country, we have designed a software tool, Fragmentation Tool_Version-1.0, for fragmentation of nucleotide sequences, (Supplementary Tool. 1). This tool helped us to fragment the SARS-CoV-2 genome sequences into 5000 base sections. We have used MultAlin online tool (18) for alignment of fragmented genome sequences with the reference viral strain sequence mentioned above. We have taken care of not to miss part of the genome sequences by having 100 bases long overlapping sequences in each fragment. We considered variation as true mutation if variations from reference sequence were identified in more than one instances.

The same practice was applied with the genome sequence samples of all six countries (Workflow: Step B). The results were validated by the MiniMap2 too as well as with published data.

### The Occurrence of 11085TTT Insertion and C27945T Mutations in the SARS-CoV-2 Genome

We downloaded all the SARS-CoV-2 genome sequences from GISAID database available on September 2, 2020, that provided us 93,265 genome sequences. We focussed our analysis for occurrence of 11085TTT insertion that translate to Phenylalanine (F) amino acid at the position 38 in non-structural protein 6 (NSP6) and C27945T mutation, which translate to stop codon after 17 amino acid in ORF-8 only from these 93,265 genome sequences (Workflow: Step C).

## Results

### Workflow of Mutation Analysis

We have used following workflow for the mutation analysis.

**Figure 1:**
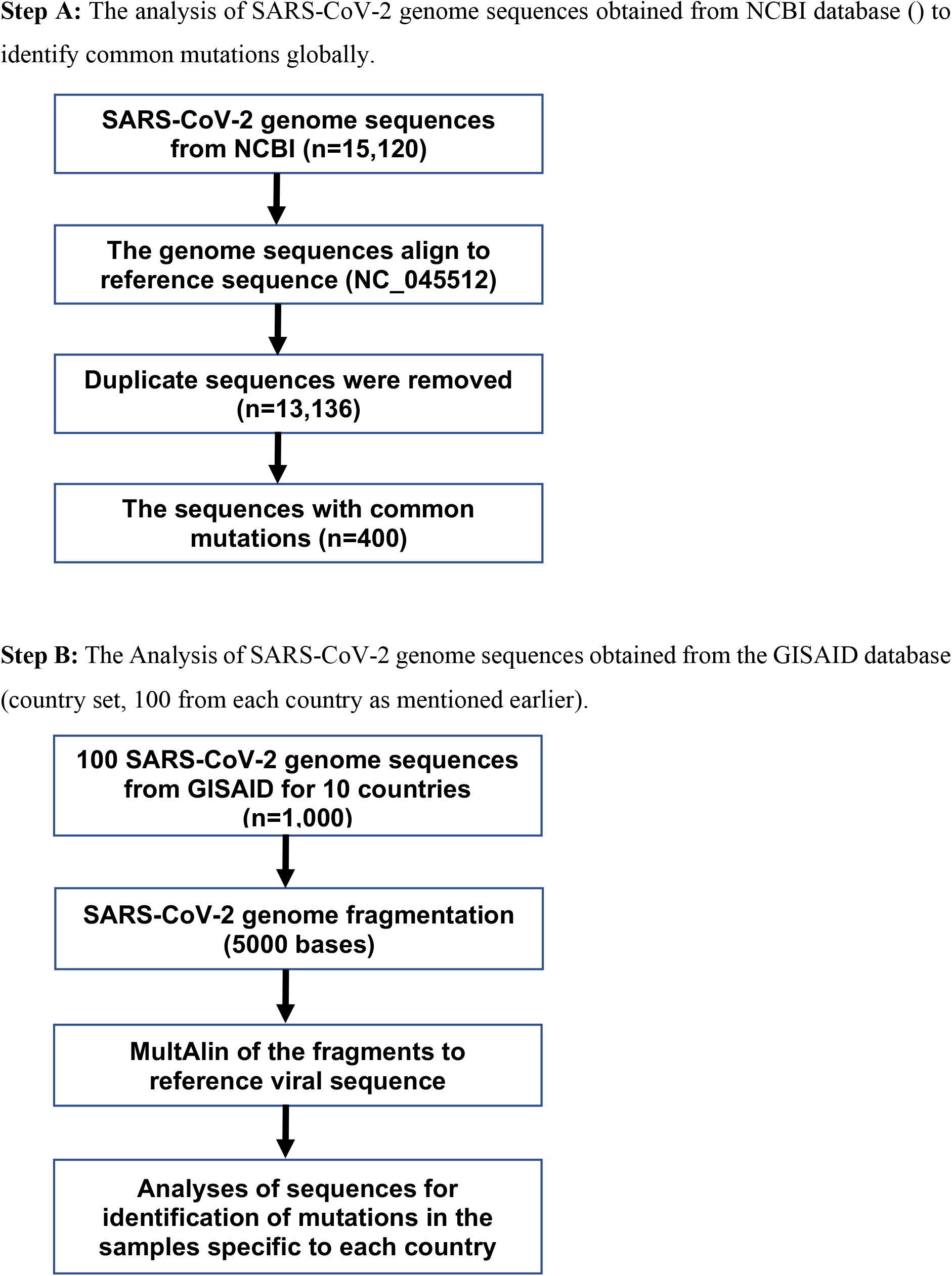
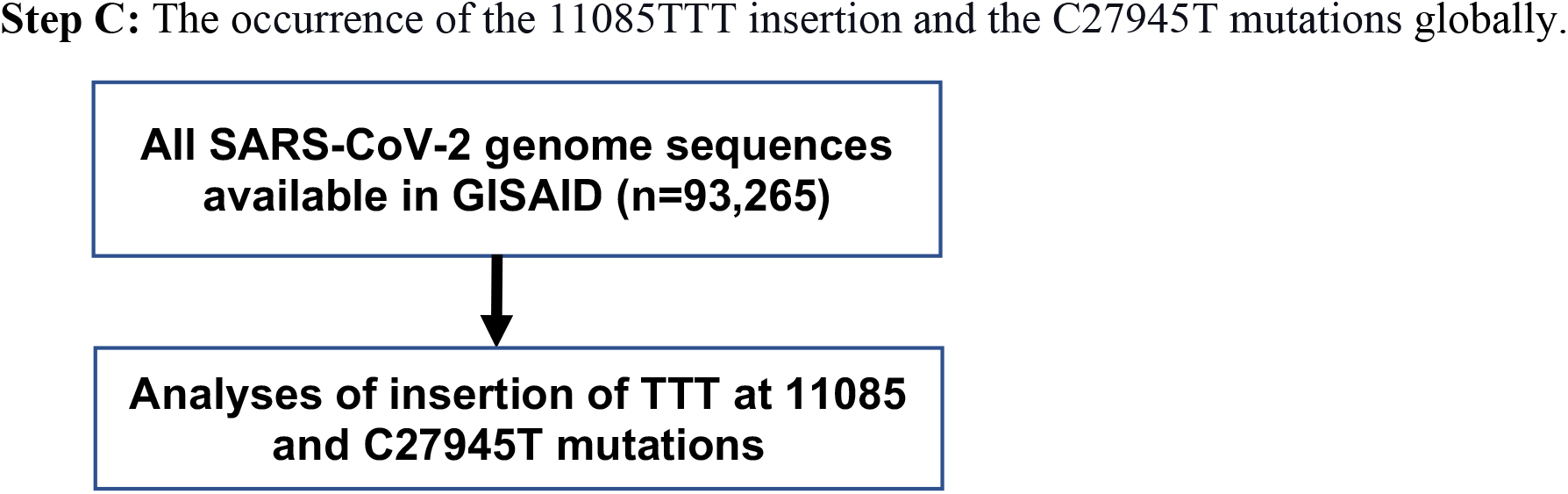
Workflow of identification of mutations in SARS-CoV-2 genome sequences.

### Common SARS-CoV-2 Mutations Across the Globe

We analysed total 15,120 full-length SARS-CoV-2 genome sequences (global set), available on the NCBI database as on 25 June 2020, investigating mutations occurrence since COVID-19 outbreak. The alignment was made with the viral reference strain (accession number NC_045512) of the virus. We detected 11 synonymous and non-synonymous mutations form the global set of data (Table 1 and Supplementary Figure 1), these mutations were previously reported (5-8, 10). From these 11 mutations the mutation C241T is occurring in 5’-UTR region, which does not get translated and so the mutations might have a regulatory role to play. The mutation was identified in quite significant number (over 85% except China USA and Spain where it is 3%, 40% and 56%, respectively) of samples from every continents. Three of the 11 mutations are synonymous mutations. The remaining 7 lead to protein change in open reading frames 1a, 1b, S, 3a and 8 (Table 1).

**Table 1:**
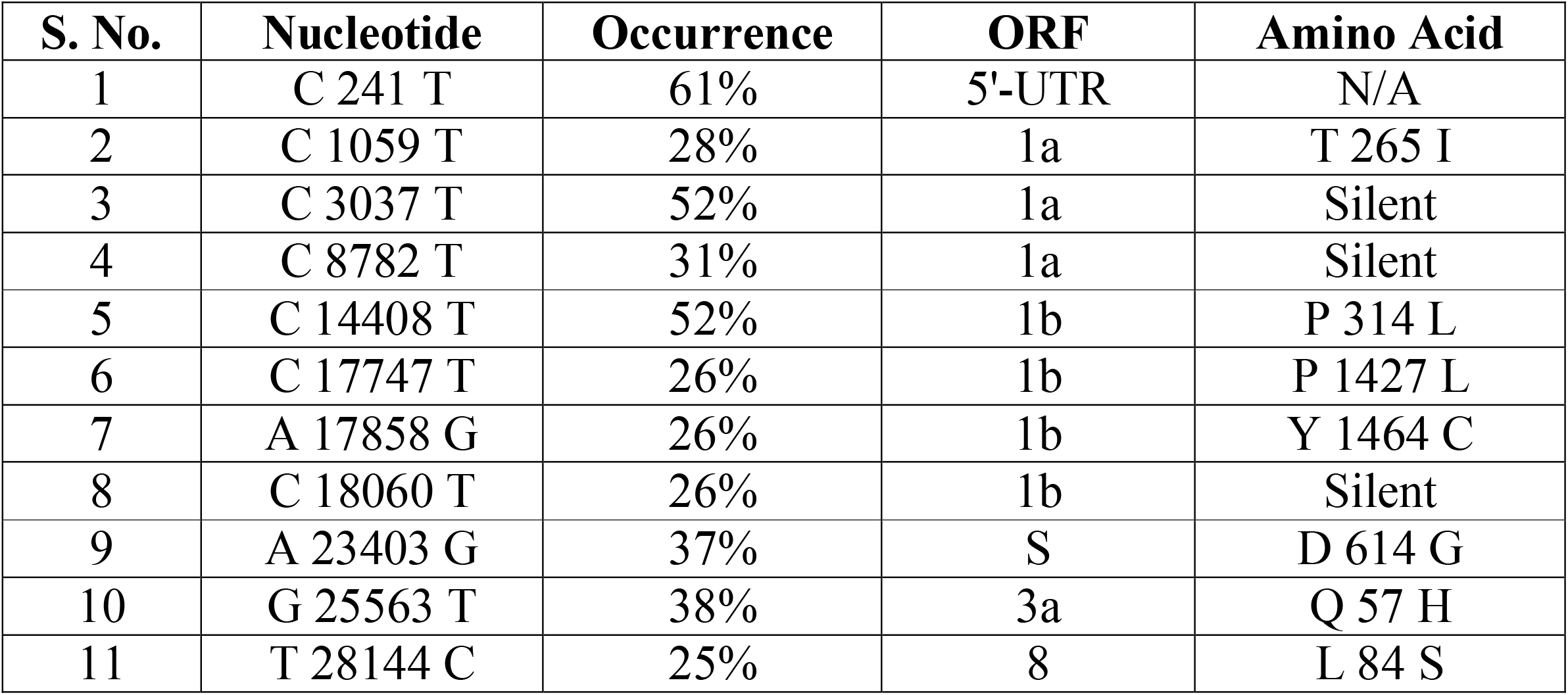
Common SARS-CoV-2 Mutations Across the Globe.

### Unique Mutation in the SARS-CoV-2 Samples from Each Continent

We were also interested in mutations occurrence in a minute percentage of samples globally. To achieve this, we have chosen Australia as representative of Oceania continent, China, India and KSA as representative of Asia, Egypt representing Africa, Italy and Spain as representative of Europe, USA for North America, Mexico for Central America and Brazil as a representative of South American continent. The chosen countries were having quite notable numbers of SARS-CoV-2 infection cases. We used the GISAID database to download the 100 full-length genome sequences derived from every above-mentioned country, selected at random. Alignment of these samples to reference strain showed us total 313 mutations in SARS-CoV-2 genome from country set of data (Supplementary Table). Many of these mutations are rare but a considerable number of mutations are common in samples derived from all six continents (Table 2). All the 11 mutations found in the global set of data were also included in the 313 mutations, obtained from the country set of data.

**Table 2:**
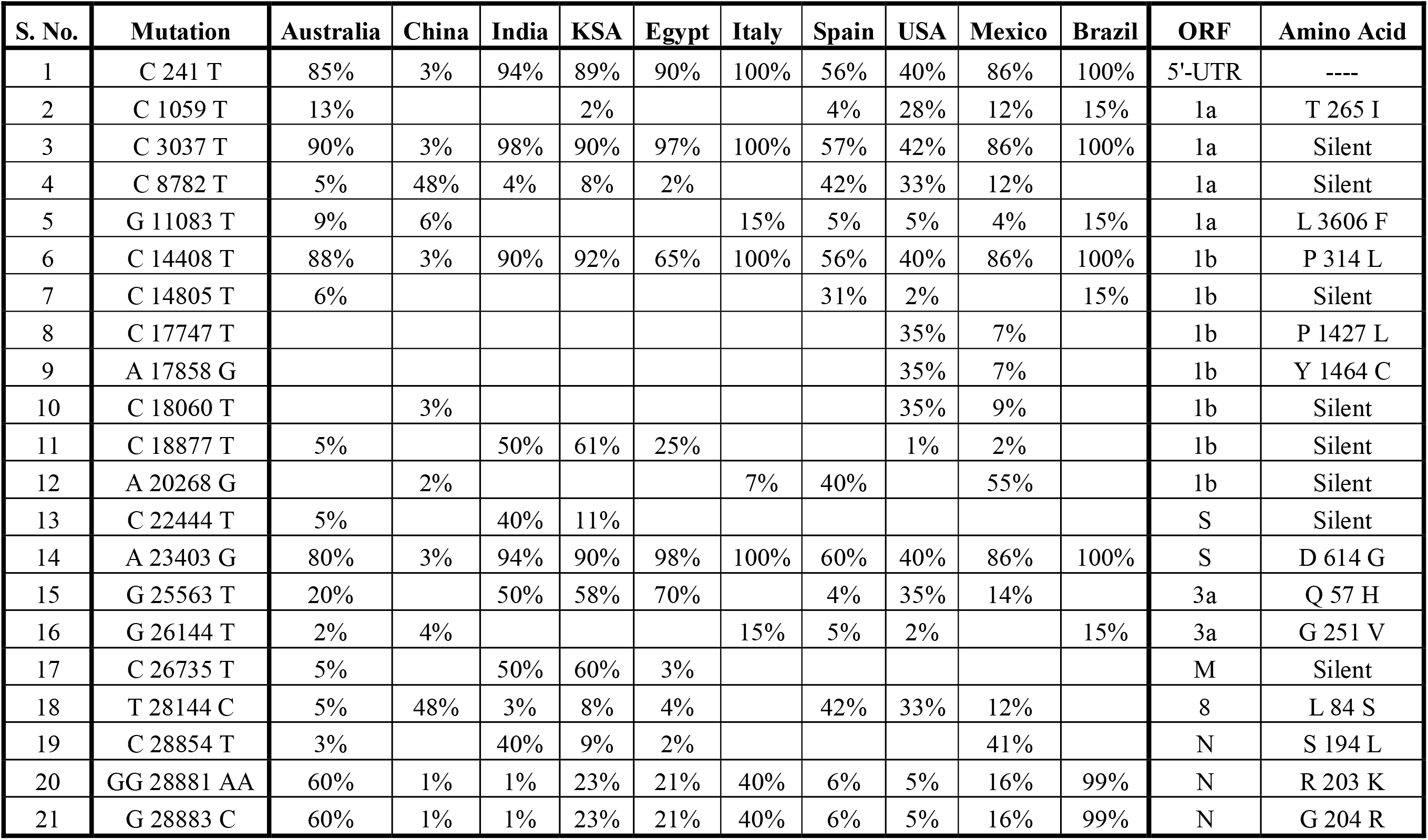
The prevalence of mutations in samples derived from representative countries of all six continents.

| S. No. | Mutation | Australia | China | India | KSA | Egypt | Italy | Spain | USA | Mexico | Brazil | ORF | Amino Acid |
| --- | --- | --- | --- | --- | --- | --- | --- | --- | --- | --- | --- | --- | --- |
| 1 | C 241 T | 85% | 3% | 94% | 89% | 90% | 100% | 56% | 40% | 86% | 100% | 5'-UTR | ---- |
| 2 | C 1059 T | 13% |  |  | 2% |  |  | 4% | 28% | 12% | 15% | 1a | T 265 I |
| 3 | C 3037 T | 90% | 3% | 98% | 90% | 97% | 100% | 57% | 42% | 86% | 100% | 1a | Silent |
| 4 | C 8782 T | 5% | 48% | 4% | 8% | 2% |  | 42% | 33% | 12% |  | 1a | Silent |
| 5 | G 11083 T | 9% | 6% |  |  |  | 15% | 5% | 5% | 4% | 15% | 1a | L 3606 F |
| 6 | C 14408 T | 88% | 3% | 90% | 92% | 65% | 100% | 56% | 40% | 86% | 100% | 1b | P 314 L |
| 7 | C 14805 T | 6% |  |  |  |  |  | 31% | 2% |  | 15% | 1b | Silent |
| 8 | C 17747 T |  |  |  |  |  |  |  | 35% | 7% |  | 1b | P 1427 L |
| 9 | A 17858 G |  |  |  |  |  |  |  | 35% | 7% |  | 1b | Y 1464 C |
| 10 | C 18060 T |  | 3% |  |  |  |  |  | 35% | 9% |  | 1b | Silent |
| 11 | C 18877 T | 5% |  | 50% | 61% | 25% |  |  | 1% | 2% |  | 1b | Silent |
| 12 | A 20268 G |  | 2% |  |  |  | 7% | 40% |  | 55% |  | 1b | Silent |
| 13 | C 22444 T | 5% |  | 40% | 11% |  |  |  |  |  |  | S | Silent |
| 14 | A 23403 G | 80% | 3% | 94% | 90% | 98% | 100% | 60% | 40% | 86% | 100% | S | D 614 G |
| 15 | G 25563 T | 20% |  | 50% | 58% | 70% |  | 4% | 35% | 14% |  | 3a | Q 57 H |
| 16 | G 26144 T | 2% | 4% |  |  |  | 15% | 5% | 2% |  | 15% | 3a | G 251 V |
| 17 | C 26735 T | 5% |  | 50% | 60% | 3% |  |  |  |  |  | M | Silent |
| 18 | T 28144 C | 5% | 48% | 3% | 8% | 4% |  | 42% | 33% | 12% |  | 8 | L 84 S |
| 19 | C 28854 T | 3% |  | 40% | 9% | 2% |  |  |  | 41% |  | N | S 194 L |
| 20 | GG 28881 AA | 60% | 1% | 1% | 23% | 21% | 40% | 6% | 5% | 16% | 99% | N | R 203 K |
| 21 | G 28883 C | 60% | 1% | 1% | 23% | 21% | 40% | 6% | 5% | 16% | 99% | N | G 204 R |

### Country Specific Mutations

In the analysis of 1000 samples, derived from representative countries of all six continents, we detected 313 total mutations, as mentioned earlier, at the nucleotide level in most of the ORFs, except in ORF 7b, 10 and 3’-UTR, (Figure 2). The highest mutations (78 mutations; 24.9%) were detected in Australian samples, (Figure 2 and Supplementary Table 1). While the lowest mutations compared to reference strain, were found in Brazilian samples (28 mutations; 8.9%) (Figure 2 and Supplementary Table 1).

**Figure 2:**
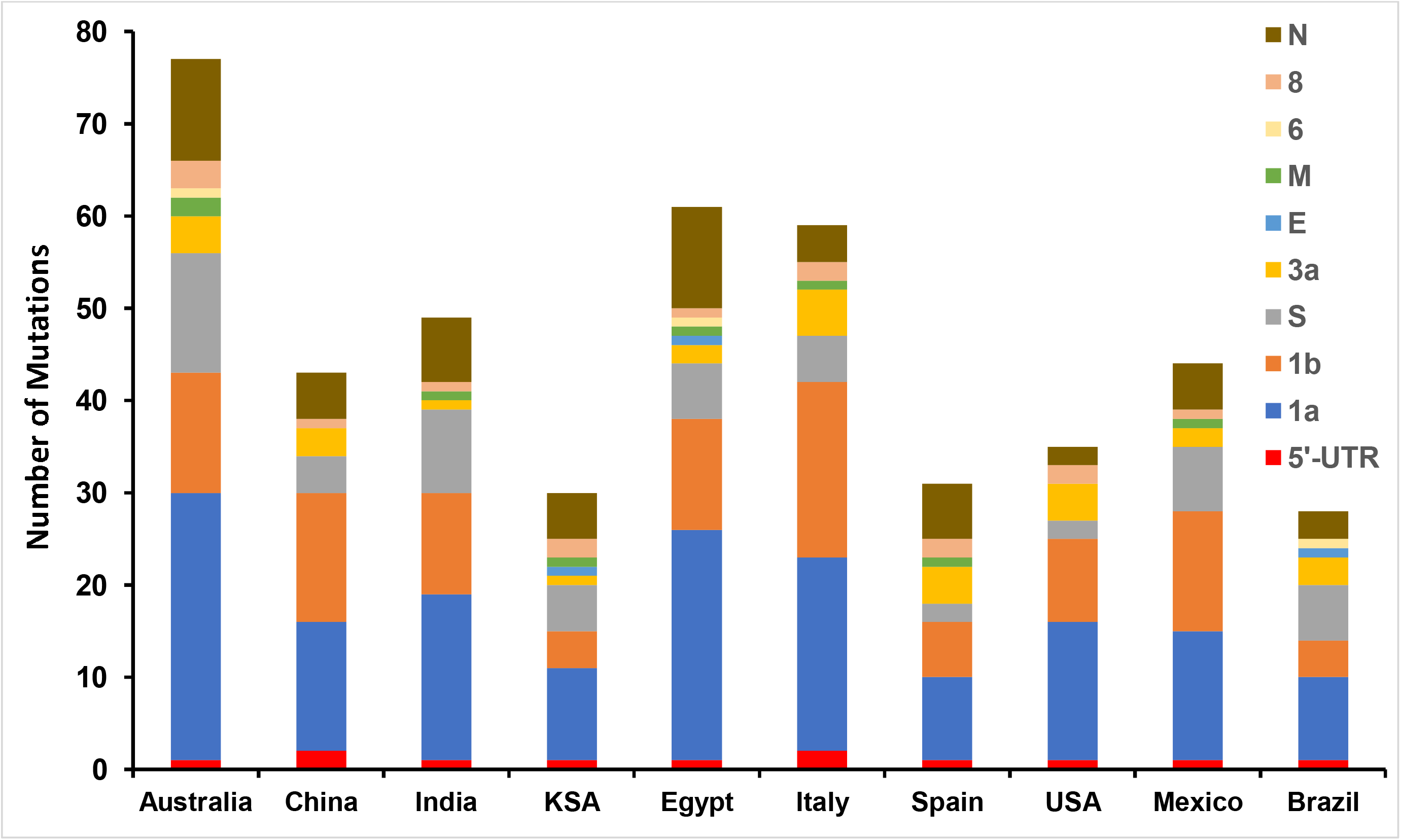
SARS-CoV-2 Mutations in different ORFs.

### Mutations at Amino Acid Level

Next, we analysed all 313 mutations, for their corresponding amino acid changes, shown in (Figure 3). The highest (43 mutations; 13.7%) non-synonymous mutations, we detected in Australian samples, which corresponds to the highest (78 mutations) nucleotide mutations. While the lowest (17 mutations; 5.4%) non-synonymous mutations compared to reference strain, were found in KSA derived samples.

**Figure 3:**
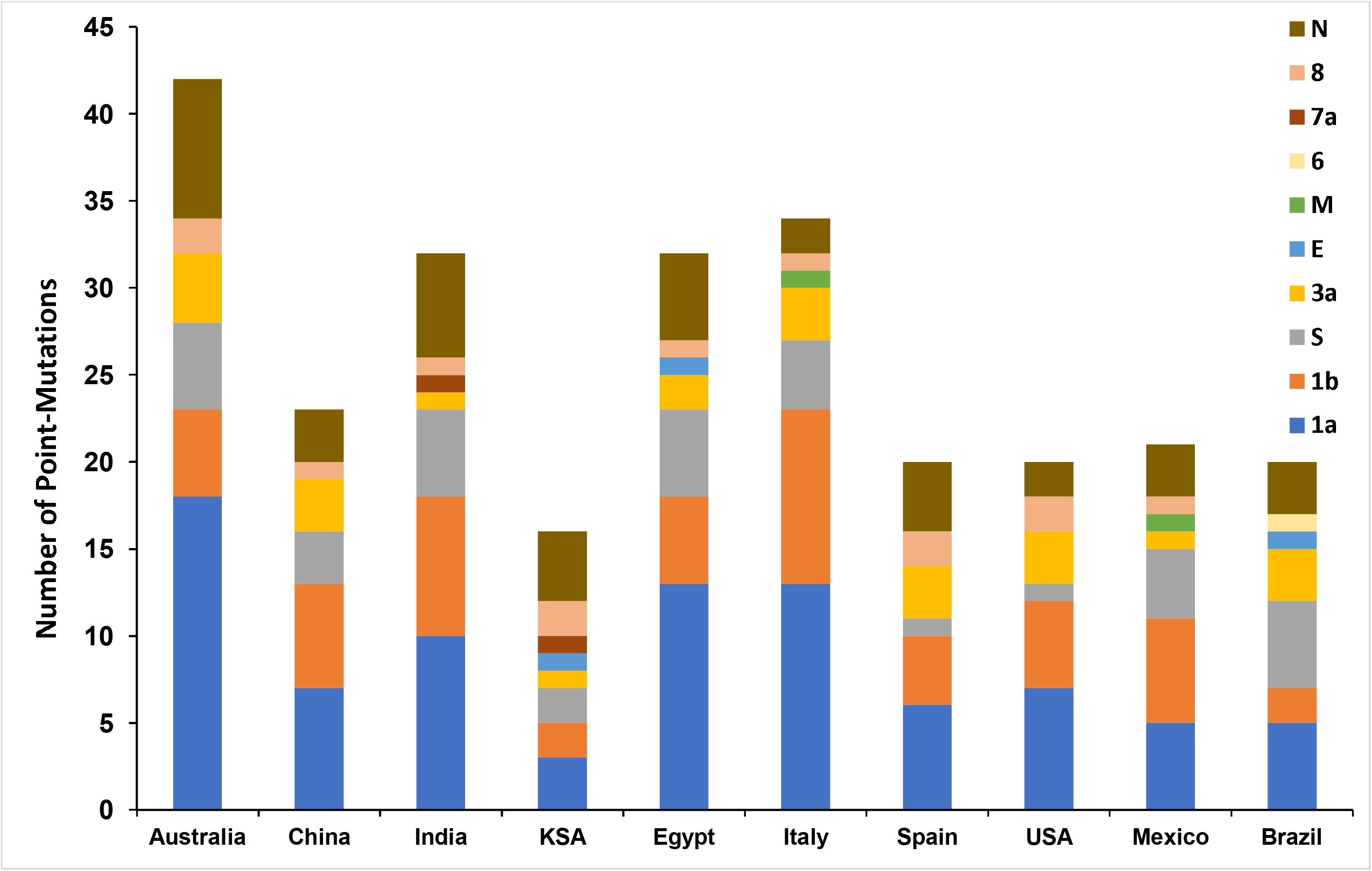
SARS-CoV-2 Non-Synonymous Mutations in different ORFs.

### Most of the Nucleotide Mutations are Non-Synonymous Mutations

We also compared non-synonymous and synonymous mutations. We noticed that non-synonymous mutations are higher than synonymous mutations in each ORF except the ORF-M, ORF-6 and ORF-N where the case was opposite (Figure 4).

**Figure 4:**
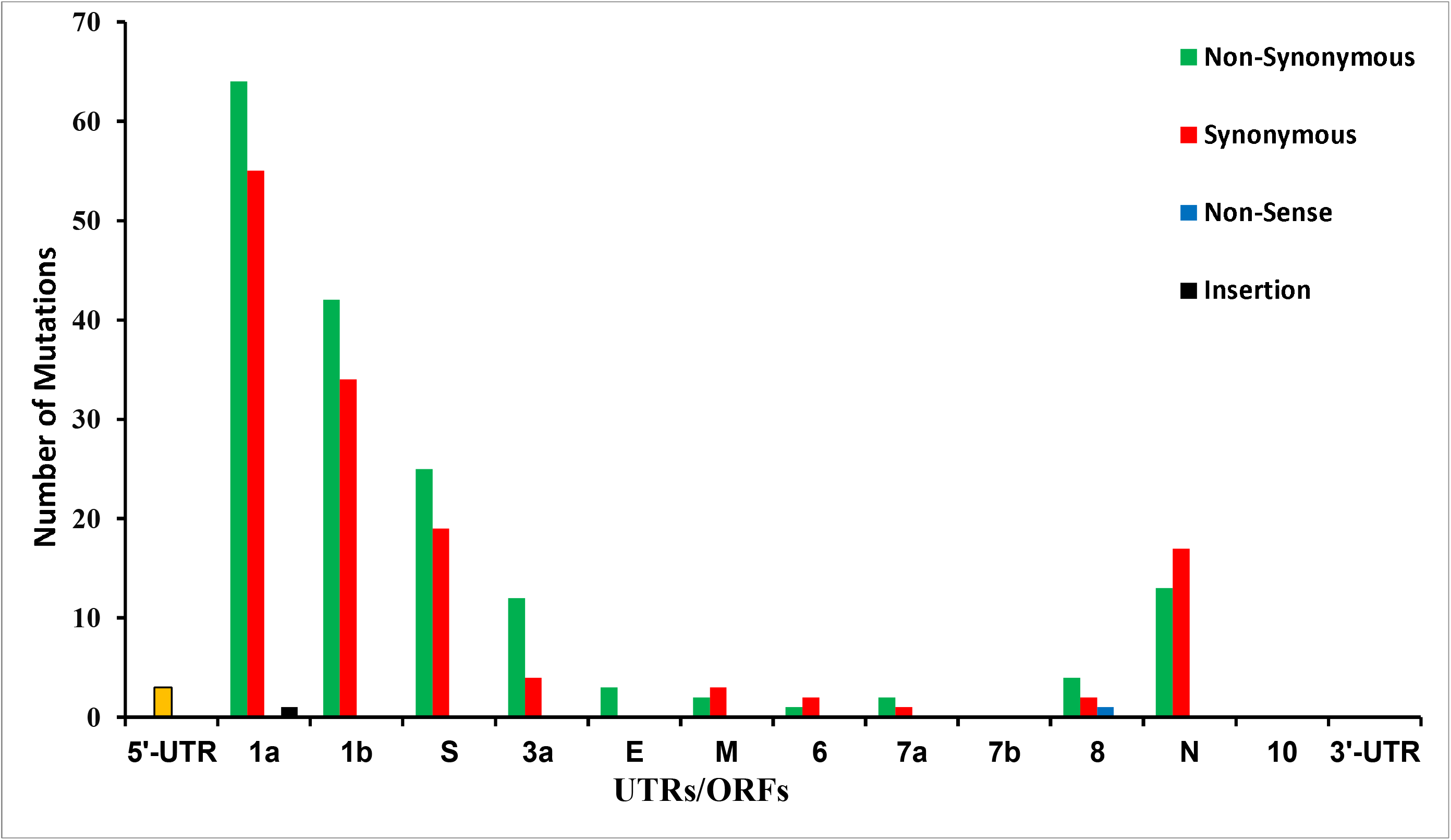
The translation of SARS-CoV-2 genomic mutation.

Interestingly, we discovered 2% of the genome sequences in Australian samples have insertion of TTT at position 11085 which lies in ORF-1a. The mutation adds one amino acid, F at the 3607 position in the ORF-1a, more precisely, the mutation adds F amino acid at the position 38 in non-structural protein 6 (NSP6), which expresses as part of ORF-1a. Another independent mutation that interests us at the position C27945T in 2% Italian samples, which translate to non-sense codon in the ORF-8. The mutation creates premature stop codon and does not allow the ORF-8 to express after the 17 amino acid although it is 121 amino acid long ORF, as per the reference strain.

### The Occurrence nsp6 Insertion Mutation Predominantly in UK Derived Samples

We analysed the occurrence of 11085TTT insertion mutation in the nsp6 in the GISAID data base. The analysis of 93,265 SARS-CoV-2 genome sequences, we found the insertion mutation in the nsp6 only in 288 samples. Remarkably, 74% of these samples were derived from the patients from UK (66%) and Australia (8%), which is the second largest after the UK (Figure 5, Supplementary Table 2).

**Figure 5:**
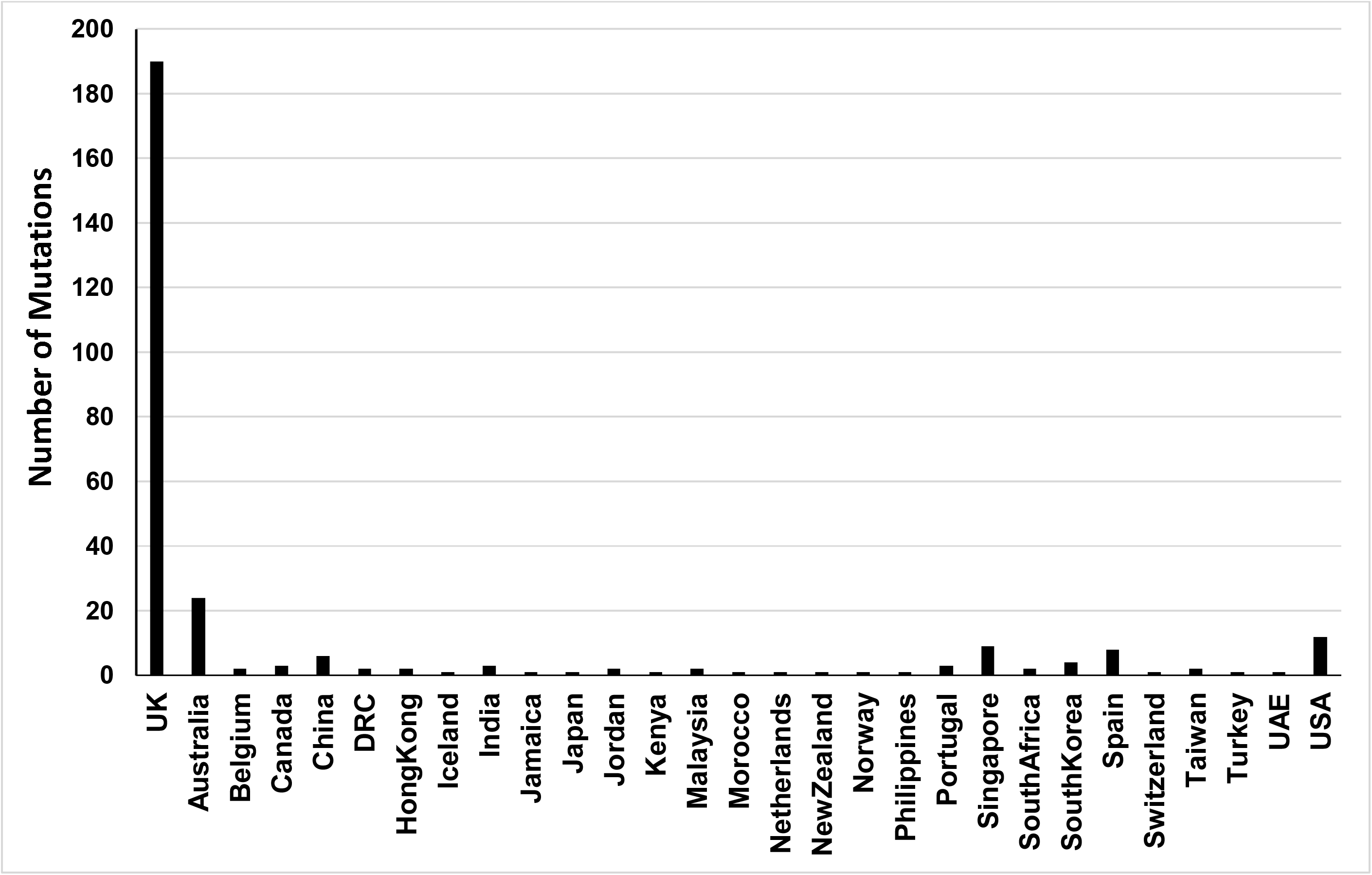
The occurrence of insertion mutation in SARS-CoV-2 nsp6 at position 38.

### The Occurrence of Non-Sense Mutation in the ORF-8 in Europe and USA Derived Samples

We also investigated the occurrence of the C27945T mutation that translate to non-sense codon after 17 amino acid in ORF-8. The search resulted to 67 samples out of 93,265 SARS-CoV-2 genome sequences obtained from the GISAID database with the C27945T mutation. 97% of these samples were derived from Europe and USA, (Figure 6A and 6B, Supplementary Figure 4 and Supplementary Table 3). Only two samples were derived from non-western countries one from Singapore and the another from South Africa.

**Figure 6:**
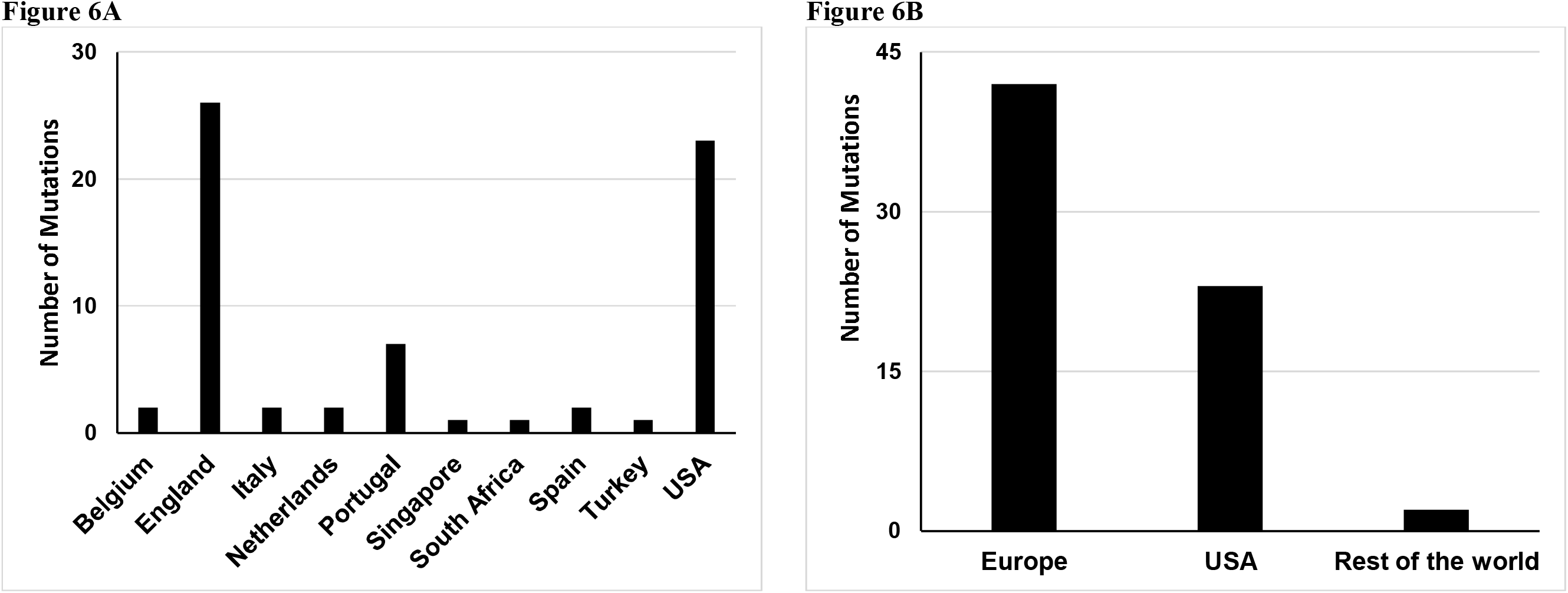
The occurrence of Non-sense mutation in SARS-CoV-2 ORF-8.

## Discussions

### Discussion

The RNA genome of SARS-CoV-2 renders the virus prone to mutations and thus changes in the UTRs and translation of the genomic code. We report novel mutations in the SARS-CoV-2 genome resulting in change in 5’-UTR or translational change, and subsequently altered proteins. The virus appears to be mutating across the globe and some of these mutations are arising independently across various populations and locations.

We detected mutations in every gene except ORF 7b, 10 and 3’-UTR. The possible reason for conserved regions in these three genes maybe the small size, or that these ORFs/UTR may not tolerate the changes which may be deleterious to the virus.

Many of the mutations we found are common in samples derived from more than one country or continent, shown in Table 2 and Supplementary Table 1. We found three mutations in 5’-UTR one of which is very common (C241T) and previously reported (8), the other two mutations occur only in 3% of samples derived from main land china and 2% of samples derived from Italy. These two independent mutations were not observed commonly in any other population, suggesting a localised non-advantageous change. The mutation C241T in 5’-UTR, was observed in 7 countries out of 10 with a frequency of more than 85%. The exceptions being China, USA and Spain where the frequency was as low as 3%, 40% and 56%, respectively. The mutation may be providing replication advantage to the virus and the mutant virus is prevailing over time. A detailed study of the C241T mutation is required to shed light on its role in infectivity or pathogenesis. Most of the identified mutations occur in ORF1ab, which is concordant with expectations of a long ORF, however, ORF-8 and ORF-N, despite their relatively small size, have a higher density of mutations as seen in Figure 4 and Supplementary table 1.

The two mutations (C17747T and A17858G) previously reported only in the USA derived samples (10), were also found in the current study with a frequency of 35% in USA samples and 7% in Mexican samples. This may be reflective of the geographical proximity of the countries and thus more frequent movement of the population. The other five mutations (C1059T, C3037T, C14408T, C18060T and A23403G), were previously reported as restricted to European derived samples (10), however, our findings indicate that these mutations are spreading in all six continents in quite significant numbers of samples over time (Table 2 and Supplementary table 1).

Although, some of the mutations are common in all samples, the frequency of majority of the mutations in all 10 countries vary. While many other mutations have different locations in the genome that indicate the virus is using a different strategy to adapt in different populations.

The A23403G mutation is translating into D614G in ORF-S, which has been observed globally (11) and correlated with mortality rate in COVID-19 patients (12). The frequency of the D614G mutation in ORF-S, was found to be more than 80% in 7 out of 10 countries, except China, USA, and Spain, where it is 3%, 40% and 60%, respectively. The close resemblance in the occurrence pattern of the mutations C241T and A23403G, two mutations at the opposite ends of the genome, in all countries is important to note, with no published record correlating the mortality rate with C241T mutation or A23403G mutation.

Out of the 313 mutations observed, the majority (169) are non-synonymous mutations. These non-synonymous mutations were found in all ORFs (except ORF 7b, 10 and 3’-UTR) and thus highly likely to affect the infection and pathogenesis of the virus profoundly (11, 12, 19).

The insertion mutation identified in NSP6 protein, which has 7 putative trans-membrane helices (20), binds to TANK binding kinase 1 (TBK1) and suppresses the phosphorylation of interferon regulatory factor 3 (IRF3) (21, 22), thereby, lowering the Type I interferon response; to evade host defences. The insertion of TTT at 11085 which adds an extra amino acid F to the NSP6 at amino acid position 38 (Supplementary Figure 2 and 3), occurs mainly in UK derived samples signifying the involvement of host’s factor in the mutation. This mutation is also present in samples from Australia (8%) and the USA (4%), further advocating the involvement of hosts possibly originating from an ancestral UK population, as Australia and USA have significant number of people originating from the UK. The plausible host genetic factors in UK originated samples need to be explored in detail, which may be responsible for the NSP6 insertion mutation.

Lastly, the C27945T mutation, translates into premature termination of ORF-8 (Supplementary Figure 4). The ORF-8 protein interacts with several host factors in the lumen of endoplasmic reticulum and also reported to be secreted out of the host’s cell (23). Unsurprisingly, it plays a role in host’s immune response evasion by disrupting the Type I interferon signalling and downregulating MHC-1 (24, 25). These observations are in concordance with published data implicating the interferon deficiency in severe COVID-19 phenotype (26). The occurrence of premature termination of ORF-8 in 97% of SARS-CoV-2 isolates derived from countries with predominantly Caucasian population, indicates the striking connection of this mutation with the said population.

## Conclusions

We identified a total of 313 synonymous and non-synonymous mutations in the SARS-CoV-2 genome across the globe. Several of these mutations were common in majority of the genome samples irrespective of geographical locations, while many other mutations were specific to a specific population. In the present study, we confirmed a link between reduced Interferon and SARS-CoV-2 mutations, and report previously unknown mutations. Two of the reported mutations appear highly linked to the location of the virus and implicate region specific mutations as a response to host environment. With the use of statistical models, such as those described by Cilia et al (27), and machine learning techniques where input from known mutations is essential, it will become possible to predict hotspots for future mutations and thus prediction of efficacy of any potential vaccines. With the emergence of new variants of SARS-CoV-2 and the possibility of any potential vaccines being rendered ineffective against the new strains, it is imperative that the structure of SARS-CoV-2 protein and genome are studied in depth in order, to enable identification of areas of the genome with a higher susceptibility to mutations.

---

**Supplementary Figure 2:**
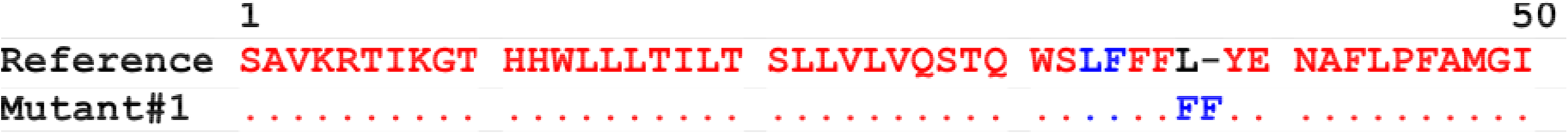
The amino acid sequence alignment of insertion mutation in nsp6 at position 38.

**Supplementary Figure 3:**
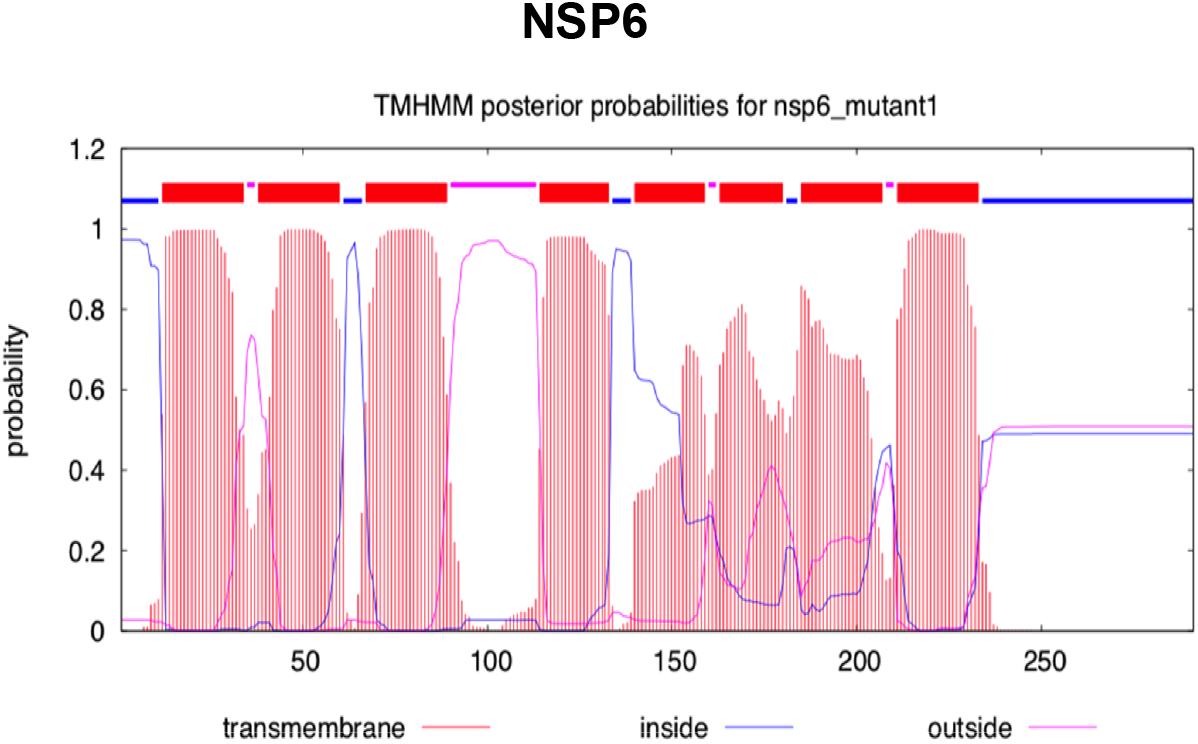
The Prediction of transmembrane domain and insertion mutation at position 38.

**Supplementary Figure 4:**
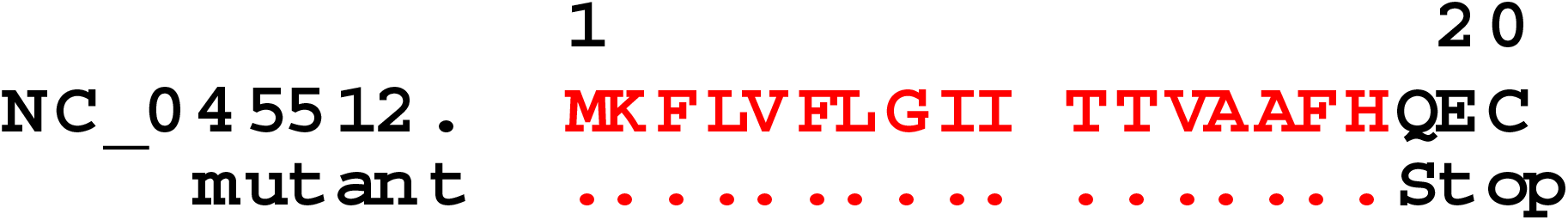
The amino acid sequence alignment of non-sense mutation in ORF-8 at position 18.

## References

1. Cui, J., Li, F., & Shi, Z. (2019). Origin and evolution of pathogenic coronaviruses. Nature Reviews Microbiology, 17(3), 181-192. doi:10.1038/s41579-018-0118-9

2. Jasper Fuk-Woo Chan, Kin-Hang Kok, Zheng Zhu, Hin Chu, Kelvin Kai-Wang To, Shuofeng Yuan and Kwok-Yung Yuen. Genomic characterization of the 2019 novel human-pathogenic coronavirus isolated from a patient with atypical pneumonia after visiting Wuhan. Emerging Microbes & Infections. 2020; 9:221–236. DOI:https://doi.org/10.1080/22221751.2020.1719902

3. Gorbalenya AE, Enjuanes L, Ziebuhr J, Snijder EJ. Nidovirales: Evolving the largest RNA virus genome. Virus Research. 2006; 117:17–37.

4. Peck KM, Lauring AS. 2018. Complexities of viral mutation rates. J Virol 92:e01031–17. https://doi.org/10.1128/JVI.01031-17.

5. Tung Phan. Genetic diversity and evolution of SARS-CoV-2. Infection, Genetics and Evolution 81 (2020) 104260. https://doi.org/10.1016/j.meegid.2020.104260

6. Tang X, Wu C, Li X, Song Y, Yao X, Wu X, Duan Y, Zhang H, Wang Y, Qian Z, Cui J, Lu J. On the origin and continuing evolution of SARS-CoV-2. National Sci Rev. 2020. https://doi.org/10.1093/nsr/nwaa036.

7. Wang C, Liu Z, Chen Z, et al. The establishment of reference sequence for SARS-CoV-2 and variation analysis. J Med Virol. 2020;92:667–674.

8. Stefanelli P. et al.. Whole genome and phylogenetic analysis of two SARSCoV-2 strains isolated in Italy in January and February 2020:Additional clues on multiple introductions and further circulation in Europe. Eurosurveillance 25, 1–5 (2020).

9. Jun-Sub Kim, Jun-Hyeong Jang, Jeong-Min Kim, Yoon-Seok Chung, Cheon-Kwon Yoo, Myung-Guk Han. Genome-Wide Identification and Characterization of Point Mutations in the SARS-CoV-2 Genome. Osong Public Health and Research Perspectives 2020; 11(3): 101–111.

10. Pachetti, M., Marini, B., Benedetti, F. et al.. Emerging SARS-CoV-2 mutation hot spots include a novel RNA-dependent-RNA polymerase variant. J Transl Med 18, 179 (2020). https://doi.org/10.1186/s12967-020-02344-6.

11. Korber, B., Fischer, W.M., Gnanakaran, S., Yoon, H., Theiler, J., Abfalterer, W., Hengartner, N., Giorgi, E.E., Bhattacharya, T., Foley, B., et al. (2020). Tracking changes in SARS-CoV-2 Spike: evidence that D614G increases infectivity of the COVID-19 virus. Cell 182, 812–827.

12. Toyoshima, Y., Nemoto, K., Matsumoto, S. et al.. SARS-CoV-2 genomic variations associated with mortality rate of COVID-19. J Hum Genet (2020). https://doi.org/10.1038/s10038-020-0808-9

13. https://www.ncbi.nlm.nih.gov/genbank/sars-cov-2-seqs/#nucleotide-sequences

14. https://www.gisaid.org

15. https://www.ncbi.nlm.nih.gov/CBBresearch/Spouge/html_ncbi/html/software/program.html?uid=11

16. Li, H. (2018). Minimap2: pairwise alignment for nucleotide sequences. Bioinformatics, 34:3094-3100. doi:10.1093/bioinformatics/bty191

17. WuF, ZhaoS, YuB, ChenYM, WangW, SongZG, etal. Anew coronavirus associated with human respiratory disease in China. Nature. 2020;579:265–9.

18. http://multalin.toulouse.inra.fr/multalin/

19. Mohammad Khalid, Hangxing Yu, Daniel Sauter, et al. (2012). “Efficient Nef-Mediated Down-modulation of TCR-CD3 and CD28 is Associated with High CD4+ T Cell Counts in Viremic HIV-2 Infection”. Journal of Virology, 2012 May; 86(9):4906–20.

20. Domenico Benvenuto, Silvia Angeletti, Marta Giovanetti, Martina Bianchi, Stefano Pascarella, Roberto Cauda, Massimo Ciccozzi, Antonio Cassone (2020). Evolutionary analysis of SARS-CoV-2: how mutation of Non-Structural Protein 6 (NSP6) could affect viral autophagy. Journal of Infection 81 (2020) e24–e27.

21. Angelini, M.M., Akhlaghpour, M., Neuman, B.W., and Buchmeier, M.J. (2013). Severe acute respiratory syndrome coronavirus non-structural proteins 3, 4, and 6 induce double-membrane vesicles. MBio 4, e00524–13.

22. Hongjie Xia, Zengguo Cao, Xuping Xie, Xianwen Zhang, John Yun-Chung Chen, Hualei Wang, Vineet D. Menachery, Ricardo Rajsbaum, and Pei-Yong Shi (2020). Evasion of Type I Interferon by SARS-CoV-2. Cell Reports 33, 108234, October 6.

23. Thomas G. Flower, Cosmo Z. Buffalo, Richard M. Hooy, Marc Allaire, Xuefeng Ren and James H. Hurley (2020). Structure of SARS-CoV-2 ORF8, a rapidly evolving coronavirus protein implicated in immune evasion. https://doi.org/10.1101/2020.08.27.270637

24. J. Y. Li et al. The ORF6, ORF8 and nucleocapsid proteins of SARS-CoV-2 inhibit type I interferon signalling pathway. Virus Res 286, 198074 (2020).

25. Y. Zhang et al., The ORF8 protein of SARS-CoV-2 mediates immune evasion through potently downregulating MHC-I. biorxiv doi.org/10.1101/2020.05.24.111823, (2020).

26. Meffre, E., and Iwasaki A. (2020). Interferon deficiency can lead to severe COVID. Nature 587, November 19, (374–375).

27. Cilia E. et al.. (2014). Predicting virus mutations through statistical relational learning. BMC Bioinformatics 15:309.

